## Supplemental information for "Translational regulation enhances distinction of cell types in the nervous system"

1: Frontier Research Institute for Interdisciplinary Sciences, Tohoku University; 2: Graduate School of Life Sciences, Tohoku University; 3: Faculty of Advanced Engineering, Tokyo University of Sciences; 4: RNA Systems Biochemistry Laboratory, RIKEN Cluster for Pioneering Research, Wako, Saitama 351-0198, Japan 5: Department of Computational Biology and Medical Sciences, Graduate School of Frontier Sciences, The University of Tokyo, Kashiwa, Chiba 277-8561, Japan

**Correspondence authors:**

Hiromu Tanimoto and Toshiharu Ichinose

**Supplementary method:**

Sequences of the biotinylated 2′-*O*-methyl oligonucleotides for rRNA-depletion.

5´-/biotin/AUUGGCAUCACAUCCAUUGUCGUUUA-3´

5´-/biotin/ACAUCCAUUGUCGUUUAUAAAGUAAA-3´

5´-/biotin/AUAAACUUUAAAUGGUUUAGAAGCCAU-3´

5´-/biotin/AAAUGGUUUAGAAGCCAUACAAUGCAAA-3´

5´-/biotin/GCCUCAUUUAAGAAGGACUUAAAUCGUUAA-3´

5´-/biotin/AACGCCCCGGGAUUGUGUUAAUUAGCUA-3´

5´-/biotin/ACGUUAUACGGGCCUGGCACCCUCUAUGGGU-3´

5´-/biotin/AUUGUUAAUCAUACAAGUGCAUAUAAU-3´

5´-/biotin/AUAAAUCUAUCAGCACUUUAUCAA-3´

5´-/biotin/UUUAAUCAAGUAAGUAAGGAAACA-3´

Sequence of the *Rh1-Venus* reporter

10xUAS-hsp70Bb_promoter-Rh1_5′ leader-Venus-Rh1_3′ UTR

TAAGCTGCAGGTCGGAGTACTGTCCTCCGAGCGGAGTACTGTCCTCCGAGCGGAGTAC
TGTCCTCCGAGCGGAGTACTGTCCTCCGAGCGGAGTACTGTCCTCCGAGCGGAGACTC
TAGCGAGCACGCGTCTGCAGGTCGGAGTACTGTCCTCCGAGCGGAGTACTGTCCTCCG
AGCGGAGTACTGTCCTCCGAGCGGAGTACTGTCCTCCGAGCGGAGTACTGTCCTCCGA
GCGGAGACTCTAGCGAGCCATATGAGCGCCGGAGTATAAATAGAGGCGCTTCGTCTAC
GGAGCGACAATTCAATTCAAACAAGCAAAACATTGCAGGTTTCCAACGACCAATCGCCG
CGACTAGTCCGCCCCAGTGAAATATTCAGAATCCAGGAACCCTTTATGTAAAAAGTGTT
AGAAATATTGTTAGTGAATTTGCAGCTTTTTATGTAGACAGTGTGATATAGGCGGATAT
AGTGACGCAGCCAGTAACCAAAACACAATGGAGAGCTTTGCAGGCGCGCCAGTGAGCA
AGGGCGAGGAGCTGTTCACCGGGGTGGTGCCCATCCTGGTCGAGCTGGACGGCGACG
TAAACGGCCACAAGTTCAGCGTGTCCGGCGAGGGCGAGGGCGATGCCACCTACGGCA
AGCTGACCCTGAAGCTCATCTGCACCACCGGCAAGCTGCCCGTGCCCTGGCCCACCCT
CGTGACCACCCTGGGCTACGGCCTGCAGTGCTTCGCCCGCTACCCCGACCACATGAA
GCAGCACGACTTCTTCAAGTCCGCCATGCCCGAAGGCTACGTCCAGGAGCGCACCATC
TTCTTCAAGGACGACGGCAACTACAAGACCCGCGCCGAGGTGAAGTTCGAGGGCGACA
CCCTGGTGAACCGCATCGAGCTGAAGGGCATCGACTTCAAGGAGGACGGCAACATCCT
GGGGCACAAGCTGGAGTACAACTACAACAGCCACAACGTCTATATCACCGCCGACAAG
CAGAAGAACGGCATCAAGGCCAACTTCAAGATCCGCCACAACATCGAGGACGGCGGC
GTGCAGCTCGCCGACCACTACCAGCAGAACACCCCCATCGGCGACGGCCCCGTGCTG

CTGCCCGACAACCACTACCTGAGCTACCAGTCCGCCCTGAGCAAAGACCCCAACGAGA
AGCGCGATCACATGGTCCTGCTGGAGTTCGTGACCGCCGCCGGGATCACTCTCGGCAT
GGACGAGCTGTACAAGTAAGCGGCCGCATTCTTTGGCGCAACAACCAGAACAGCAACA
ACAACAACAAGAACATCTAACTACTTACAACAGCAACAACAACAGCAACAAAAACAACAG
CAAGAACAACTGCAGCAACAGAACGAAACGCTTTCGAATAACATCAAAAACTTCAACAAT
AATGAAAAAATTATGCAACTTTCTTACATAACAAAAAGCAATGTAAACTCAGTTATTAAAT
TTCCTGCAATGTCAGTTAAGGACAAAAAAAAACTCAACAAAAAAAAATAAATGCAAACGAA
CTAGAAAAGTTATAAATTAAAATGAGCCTTTTCAAAACATAGTATATCTAACAAAAGCAGC
TTTTAGCGTGGAAAAACCCTAATGACGAACCTACAAAAGTTCGGATATCAACTTTCGGTT
ATCTTTCGCCTTTAAAGTTTGGAGAACCACAACAAATTTGAGTTTATTCATTCTTATATGT
ATAATAGTCTTCTTCAGAAGCTATAAATCCTTTCCAGGCATGCACTTGGCTT

Sequence of the mutated *Rh1-Venus* reporter

10xUAS-hsp70Bb_promoter-Rh1_m_5′ leader-Venus-Rh1_3′ UTR

TAAGCTGCAGGTCGGAGTACTGTCCTCCGAGCGGAGTACTGTCCTCCGAGCGGAGTAC
TGTCCTCCGAGCGGAGTACTGTCCTCCGAGCGGAGTACTGTCCTCCGAGCGGAGACTC
TAGCGAGCACGCGTCTGCAGGTCGGAGTACTGTCCTCCGAGCGGAGTACTGTCCTCCG
AGCGGAGTACTGTCCTCCGAGCGGAGTACTGTCCTCCGAGCGGAGTACTGTCCTCCGA
GCGGAGACTCTAGCGAGCCATATGAGCGCCGGAGTATAAATAGAGGCGCTTCGTCTAC
GGAGCGACAATTCAATTCAAACAAGCAAAACATTGCAGGTTTCCAACGACCAATCGCCG
CGACTAGTCCGCCCCAGTGAAATATTCAGAATCCAGGAACCCTTTCCCAAAAAAGTGTT
AGAAATATTGTTAGTGAATTTGCAGCTTTTTCCCAAAACAGTGTGATATAGGCGGGCCCA
AACGACGCAGCCAGTAACCAAAACACAATGGAGAGCTTTGCAGGCGCGCCAGTGAGCA
AGGGCGAGGAGCTGTTCACCGGGGTGGTGCCCATCCTGGTCGAGCTGGACGGCGACG
TAAACGGCCACAAGTTCAGCGTGTCCGGCGAGGGCGAGGGCGATGCCACCTACGGCA
AGCTGACCCTGAAGCTCATCTGCACCACCGGCAAGCTGCCCGTGCCCTGGCCCACCCT
CGTGACCACCCTGGGCTACGGCCTGCAGTGCTTCGCCCGCTACCCCGACCACATGAA
GCAGCACGACTTCTTCAAGTCCGCCATGCCCGAAGGCTACGTCCAGGAGCGCACCATC
TTCTTCAAGGACGACGGCAACTACAAGACCCGCGCCGAGGTGAAGTTCGAGGGCGACA
CCCTGGTGAACCGCATCGAGCTGAAGGGCATCGACTTCAAGGAGGACGGCAACATCCT
GGGGCACAAGCTGGAGTACAACTACAACAGCCACAACGTCTATATCACCGCCGACAAG
CAGAAGAACGGCATCAAGGCCAACTTCAAGATCCGCCACAACATCGAGGACGGCGGC
GTGCAGCTCGCCGACCACTACCAGCAGAACACCCCCATCGGCGACGGCCCCGTGCTG
CTGCCCGACAACCACTACCTGAGCTACCAGTCCGCCCTGAGCAAAGACCCCAACGAGA

AGCGCGATCACATGGTCCTGCTGGAGTTCGTGACCGCCGCCGGGATCACTCTCGGCATGGACGAGCTGTACAAGTAAGCGGCCGCATTCTTTGGCGCAACAACCAGAACAGCAACAACAACAACAAGAACATCTAACTACTTACAACAGCAACAACAACAGCAACAAAAACAACAGCAAGAACAACTGCAGCAACAGAACGAAACGCTTTCGAATAACATCAAAAACTTCAACAATAATGAAAAAATTATGCAACTTTCTTACATAACAAAAAGCAATGTAAACTCAGTTATTAAATTTCCTGCAATGTCAGTTAAGGACAAAAAAAAACTCAACAAAAAAAATAAATGCAAACGAACTAGAAAAGTTATAAATTAAAATGAGCCTTTTCAAAACATAGTATATCTAACAAAAGCAGCTTTTAGCGTGGAAAAACCCTAATGACGAACCTACAAAAGTTCGGATATCAACTTTCGGTTATCTTTCGCCTTTAAAGTTTGGAGAACCACAACAAATTTGAGTTTATTCATTCTTATATGTATAATAGTCTTCTTCAGAAGCTATAAATCCTTTCCAGGCATGCACTTGGCTT

Sequences of the *Venus* or *GFP* probes (18 nt, 31 probes)

#1:CGGTGAACAGCTCCTCGC, #2:GACCAGGATGGGCACCAC, #3:GTTTACGTCGCCGTCCAG, #4:CCGGACACGCTGAACTTG, #5:TTGCCGTAGGTGGCATCG, #6:GTGGTGCAGATGAGCTTC, #7:AGGGTGGTCACGAGGGTG, #8: AAGCACTGCAGGCCGTAG, #9:ATGTGGTCGGGGTAGCGG, #10:TGAAGAAGTCGTGCTGCT, #11:ACGTAGCCTTCGGGCATG, #12:AGAAGATGGTGCGCTCCT, #13:TCTTGTAGTTGCCGTCGT, #14:TCGAACTTCACCTCGGCG, #15:TCGATGCGGTTCACCAGG, #16:TGAAGTCGATGCCCTTCA, #17:CAGGATGTTGCCGTCCTC, #18:TTGTACTCCAGCTTGTGC, #19:AGACGTTGTGGCTGTTGT, #20:TTCTTCTGCTTGTCGGCG, #21:TGAAGTTGGCCTTGATGC, #22:CTCGATGTTGTGGCGGAT, #23:TAGTGGTCGGCGAGCTGC, #24:CGATGGGGGTGTTCTGCT, #25:TTGTCGGGCAGCAGCACG, #26:GGACTGGTAGCTCAGGTA, #27:GTTGGGGTCTTTGCTCAG, #28:ACCATGTGATCGCGCTTC, #29:CGGTCACGAACTCCAGCA, #30:TCCATGCCGAGAGTGATC, #31:CCGCTTACTTGTACAGCT

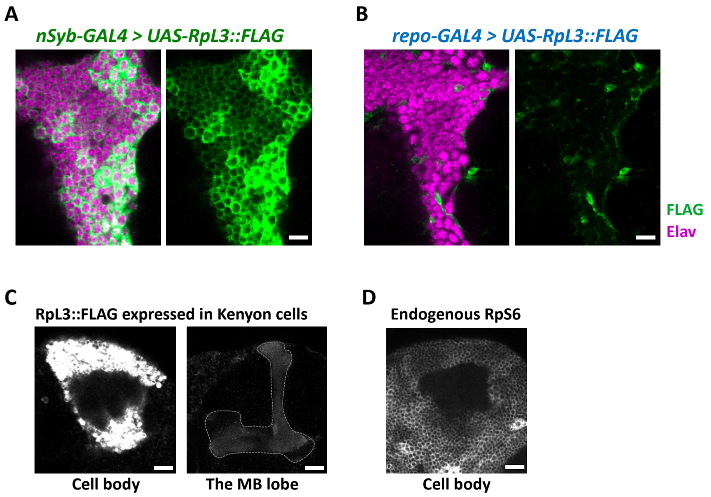


**Figure 2 – figure supplement 1. Cell-type specific ribosome profiling.**

(A-B) Immunohistochemical signal of the FLAG-tagged RpL3 protein (green) and the neuronal marker elav protein (magenta) of the indicated genotypes. Sliced confocal images of the cortical regions adjacent to the antennal lobe are shown. Scale bars: 10 µm. (C) Immunohistochemical signal of the FLAG-tagged RpL3 protein in Kenyon cells, driven by *MB010B*. The mushroom body lobe is outlined by the dotted line. Scale bars: 20 µm. (D) Immunohistochemical signal of the endogenous RpS6 protein in the wild type brain. Cortical region of the posterior side containing Kenyon cells are shown. Scale bar: 20 µm.

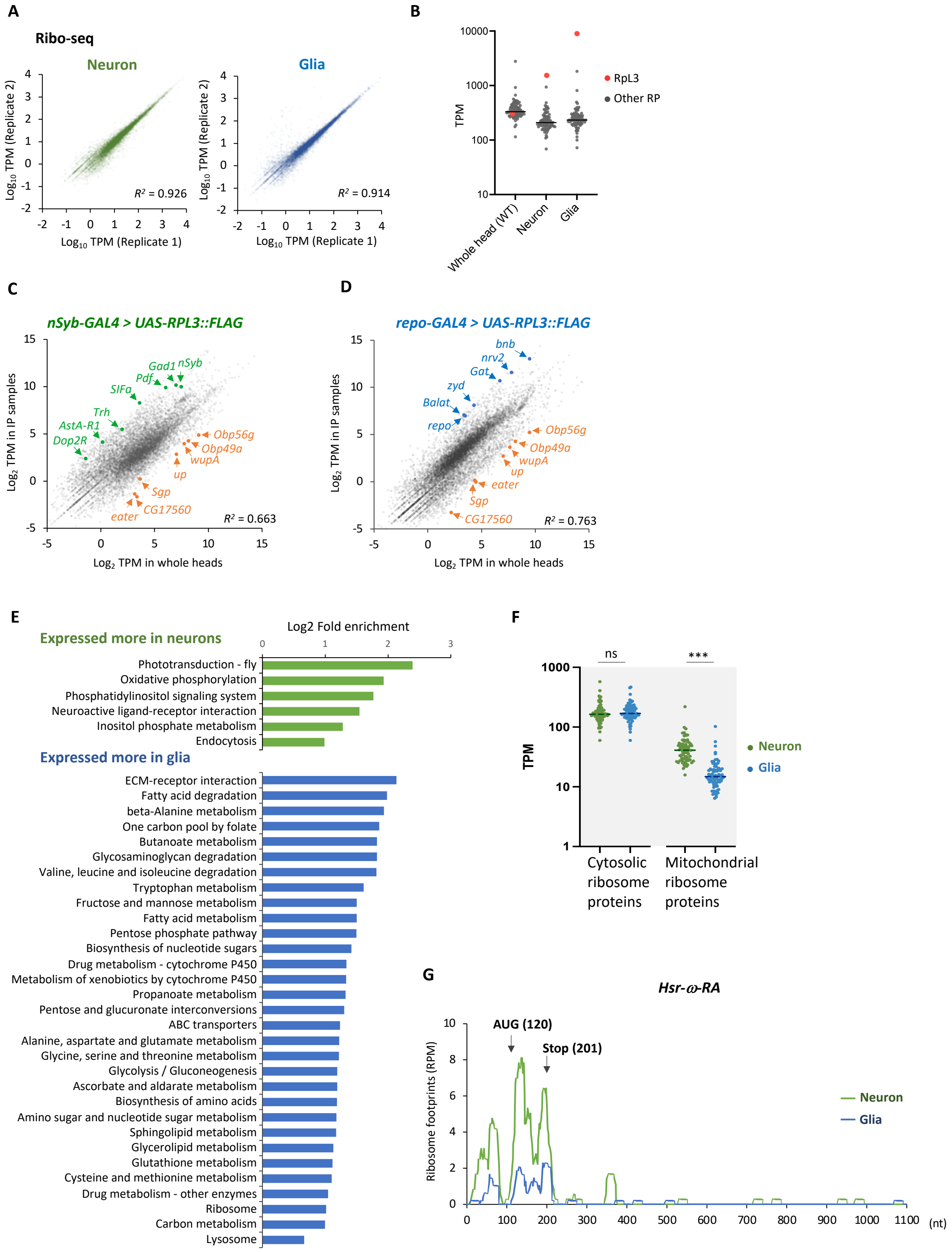
A
Ribo-seq
Neuron
Glia
Log10 TPM (Replicate 2)
Log10 TPM (Replicate 1)
R² = 0.926
R² = 0.914
B
TPM
RpL3
Other RP
Whole head (WT)
Neuron
Glia
C
nSyb-GAL4 > UAS-RPL3::FLAG
Gad1
nSyb
Pdf
SIFa
Trh
AstA-R1
Dop2R
Obp56g
Obp49a
wupA
up
Sgp
CG17560
eater
R² = 0.663
Log2 TPM in IP samples
Log2 TPM in whole heads
D
repo-GAL4 > UAS-RPL3::FLAG
bnb
nrv2
Gat
zyd
Balat
repo
Obp56g
Obp49a
wupA
up
eater
Sgp
CG17560
R² = 0.763
E
Log2 Fold enrichment
Expressed more in neurons
Phototransduction - fly
Oxidative phosphorylation
Phosphatidylinositol signaling system
Neuroactive ligand-receptor interaction
Inositol phosphate metabolism
Endocytosis
Expressed more in glia
ECM-receptor interaction
Fatty acid degradation
beta-Alanine metabolism
One carbon pool by folate
Butanoate metabolism
Glycosaminoglycan degradation
Valine, leucine and isoleucine degradation
Tryptophan metabolism
Fructose and mannose metabolism
Fatty acid metabolism
Pentose phosphate pathway
Biosynthesis of nucleotide sugars
Drug metabolism - cytochrome P450
Metabolism of xenobiotics by cytochrome P450
Propanoate metabolism
Pentose and glucuronate interconversions
ABC transporters
Alanine, aspartate and glutamate metabolism
Glycine, serine and threonine metabolism
Glycolysis / Gluconeogenesis
Ascorbate and aldarate metabolism
Biosynthesis of amino acids
Amino sugar and nucleotide sugar metabolism
Sphingolipid metabolism
Glycerolipid metabolism
Glutathione metabolism
Cysteine and methionine metabolism
Drug metabolism - other enzymes
Ribosome
Carbon metabolism
Lysosome
F
ns
***
TPM
Neuron
Glia
Cytosolic ribosome proteins
Mitochondrial ribosome proteins
G
Hsr-ω-RA
AUG (120)
Stop (201)
Ribosome footprints (RPM)
Neuron
Glia
(nt)

**Figure 2 – figure supplement 2. Cell-type specific ribosome profiling.**

(A) Correlation plots among the biological replicates in the cell-type specific ribo-seq (Transcripts per million; TPM). The squared Pearson's correlation coefficient ($R^2$) is indicated. (B) Reads on the cytosolic ribosome proteins in the whole head sample (wild type), neurons, or the glial cells. The red and grey points represent the TPM of RpL3 and the other ribosome proteins, respectively. Note that RpL3::FLAG is overexpressed in the neuron and the glia samples so that the reads are from both the endogenous and exogenous RpL3. (C-D) Ribosome footprints in immunoprecipitated (y-axis) and the whole head (x-axis) samples from the indicated genotypes are plotted (TPM). Enrichment of neuronal (green) or glial (blue) marker genes and depletion of markers for other cell types (orange), including muscles (*wupA* and *up*), fat bodies (*Sgp* and *CG17560*), hemocytes (*eater*), and auxiliary cells (*Obp56g* and *Obp49a*), are highlighted. The squared Pearson's correlation coefficient ($R^2$) is indicated. (E) Kyoto Encyclopedia of Genes and Genomes (KEGG) pathways enrichment analysis on differentially expressed genes among neuronal and glial cells. 1,853 or 1,657 genes are expressed significantly more in neurons or in glia, respectively (DEseq, $FDR < 0.05$), and the enrichment is tested using Database for Annotation, Visualization, and Discovery (DAVID) (Dennis et al., 2003). KEGG pathways showing $P$ values smaller than 0.005 are shown. (F) Ribosome footprints (TPM) on CDS of cytosolic or mitochondrial ribosome proteins in neurons or in glia. ns: $P > 0.05$, ***: $P < 0.001$, Mann-Whitney test of ranks. (G) Ribosome footprints on *Hsr-ω-RA* (*CR31400*) in neurons (green) or in glia (blue). A putative ORF, consisting of 81 bases, is indicated with arrows.

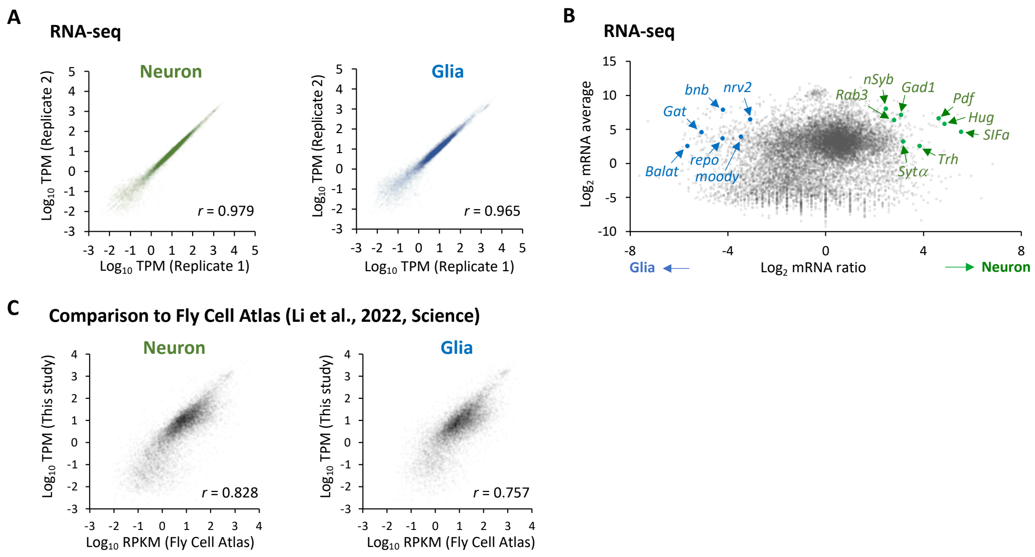


**Figure 2 – figure supplement 3. Cell-type specific RNA-seq.**

(A) Correlation plots among the biological replicates in the cell type specific RNA-seq. The squared Pearson's correlation coefficient (*r*) is indicated. (B) The MA-plot of RNA-seq from neurons and glia. Each transcript is plotted based on the fold change (x-axis) and the average (y-axis) in the unit of $\log_2$. Several marker genes are highlighted with green (neuron) or blue (glia).

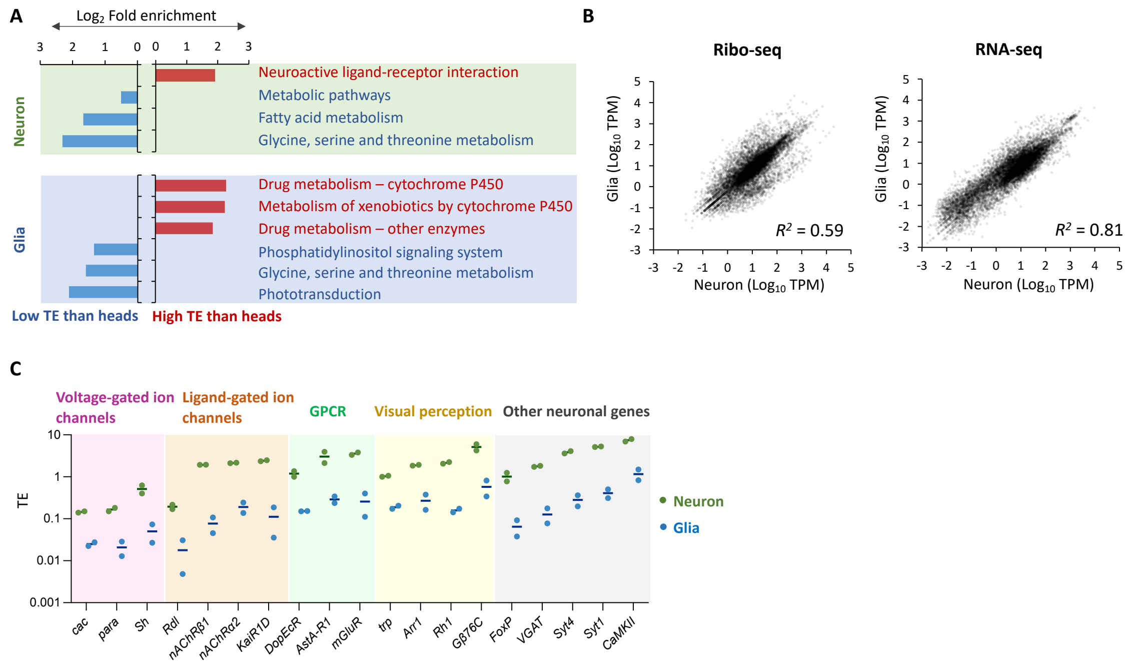


**Figure 2 – figure supplement 4. Translational efficiency in neurons, glial cells and the whole heads.**

(A) Fold enrichment of genes that are translationally enhanced or suppressed in neurons (top) or in glia (bottom), compared to the whole heads. Genes with TPM > 1 in RNA-seq are categorized as translationally enhanced if their TE in neurons or in glia is more than twice the TE in the whole head, or as suppressed if their TE in neurons or in glia is less than half of that in the whole head. KEGG enrichment analysis is performed using Database for Annotation, Visualization, and Discovery (DAVID) (Dennis et al., 2003). KEGG pathways with $P < 0.005$ are shown. (B) Correlation plot among neurons (x-axis) and glia (y-axis), with the RNA-seq read counts (left, TPM) and the ribo-seq read counts on CDS (right, TPM). The squared Pearson's correlation coefficient ($R^2$) is indicated. (C) Translational efficiency of representative neuronal genes in the indicated GO terms shown in Figure 2G-H (green: neurons; blue: glia). The bars represent the mean of the two biological replicates.

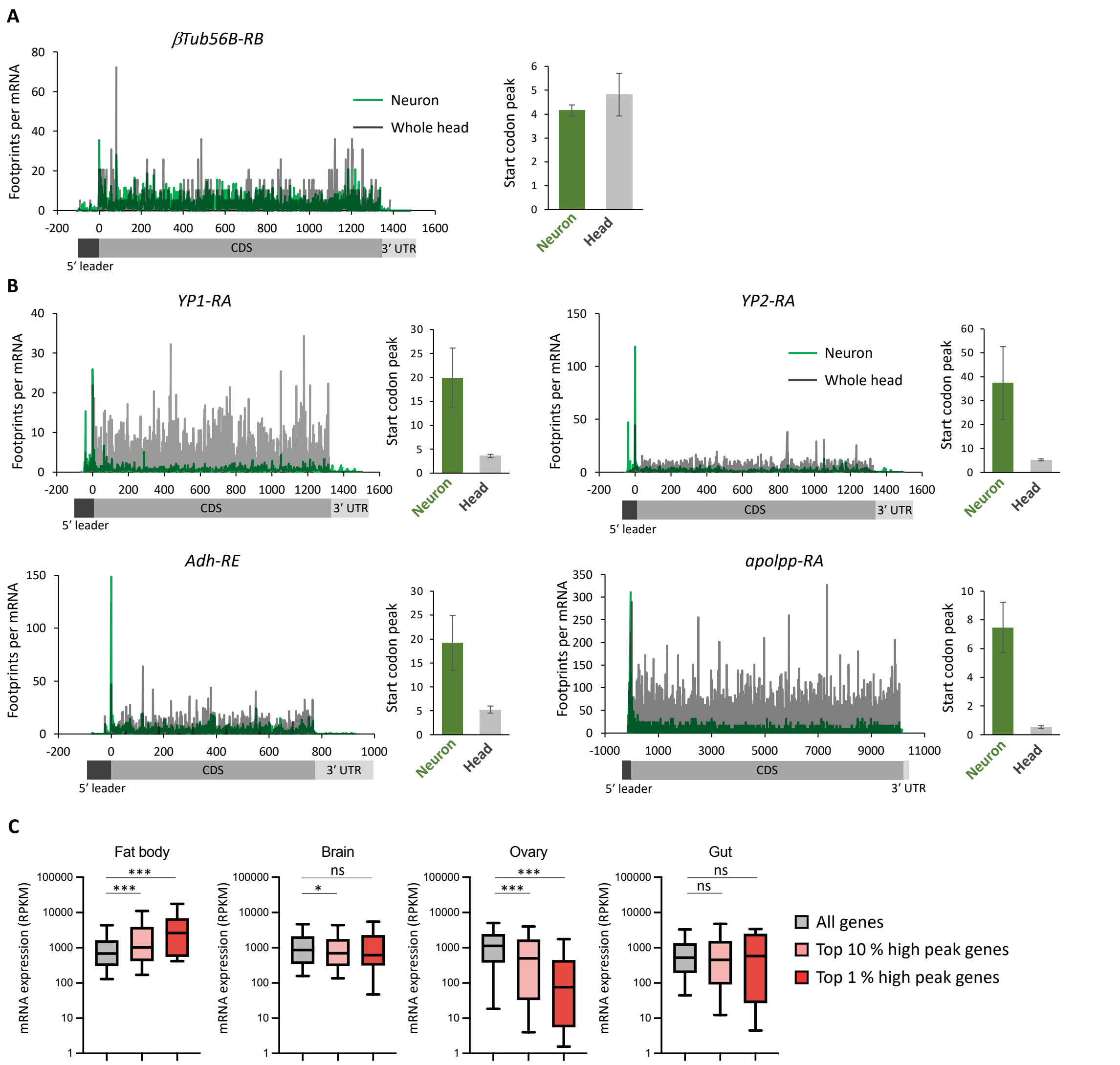


**Figure 3 – figure supplement 1. Ribosome stall at the initiation codons in neurons.**

(A-B) Ribosome distribution (P-sites, RPM), normalized by the mRNA level (TPM), on a house keeping gene, *β-tubulin at 56D* (A) or genes highly expressed in fat bodies (B) (Dobson et al., 2018). Green and grey lines indicate the distribution in neurons and the whole head, respectively. Start codon peak, calculated as footprints around the start codon (TPM, ± 2 bases) normalized by the TPM in the whole coding sequence, is plotted. Bars and error bars: mean and standard error of mean. (C) All genes showing TPM > 5 (Ribo-seq, CDS) are grouped according to the rank of start codon peak in neurons (top 10 % and 1 %). mRNA level in the indicated tissues, measured in (Dobson et al., 2018), is plotted (box and whiskers represent 25-75 and 5-95 percentile, respectively). ns: $P > 0.05$, *: $P < 0.05$, ***: $P < 0.001$, Dunn's multiple comparisons test.

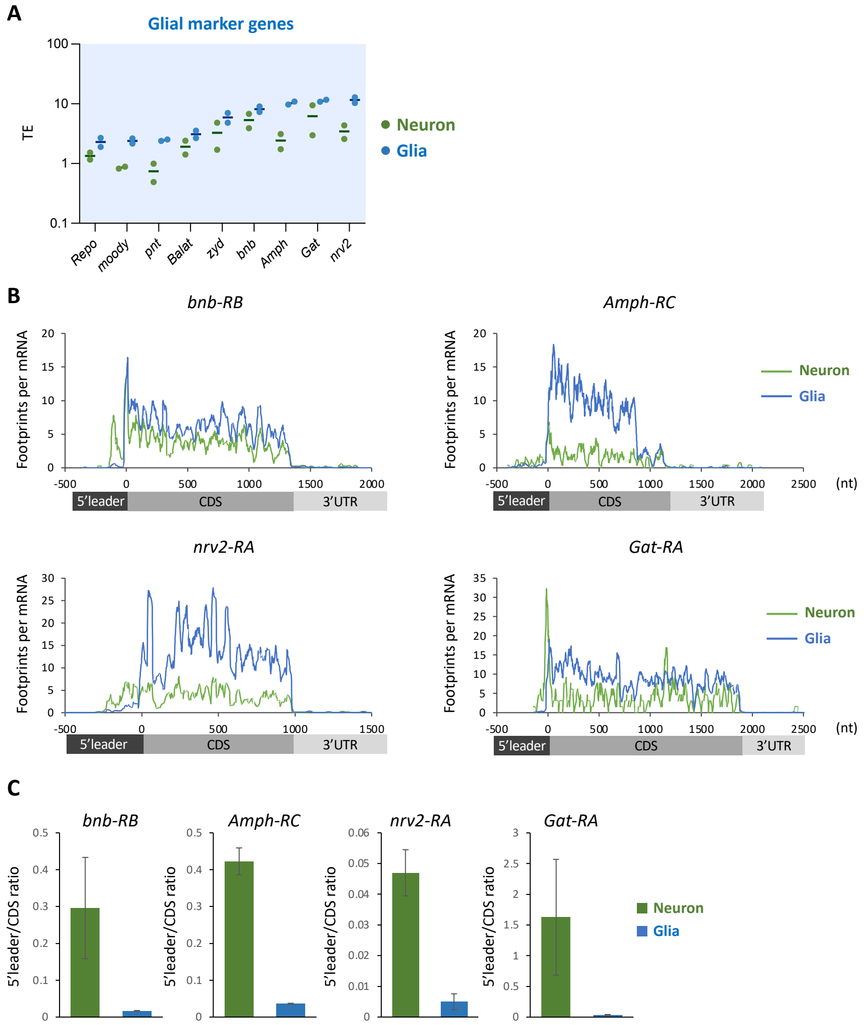


**Figure 3 – figure supplement 2. Translational suppression of glial marker genes in neurons.**

(A) Translational efficiency (TE) of the major glial marker genes in neurons (green) or in glia (blue). The bars represent the mean of the two biological replicates. (B) Ribosome distribution (P-sites, RPM), normalized by the mRNA level (TPM). Green and grey lines indicate the distribution in neurons and the whole head, respectively. Green and blue lines indicate the distribution in neuronal and glial cells, respectively. (C) Ratio of ribosome density on 5′ leader to CDS (mean ± standard error of mean of the biological replicates).

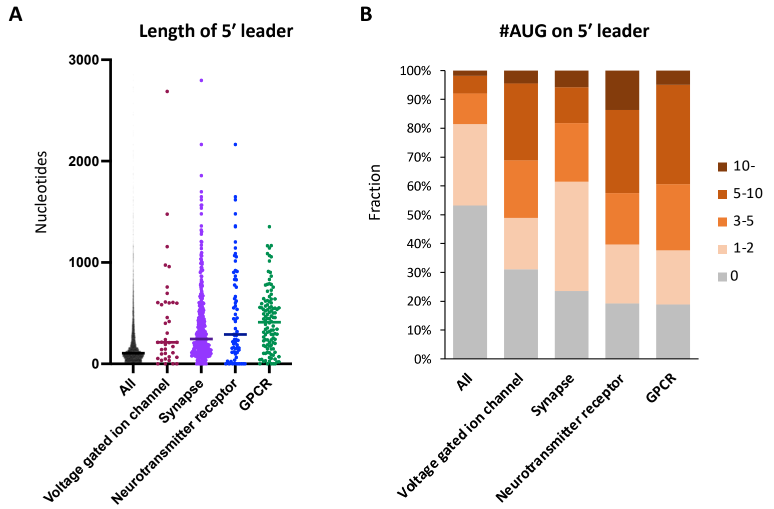


**Figure 4 – figure supplement 1. Neuronal transcripts harbor long 5′ UTR containing numerous uORF.**

(A) Length of 5′ UTR of the transcripts in the indicated Gene Ontology (GO) terms. The bars represent the median length. (B) Proportion of transcripts in the indicated GO terms, based on the number of upstream AUG codons in their 5′ UTR.

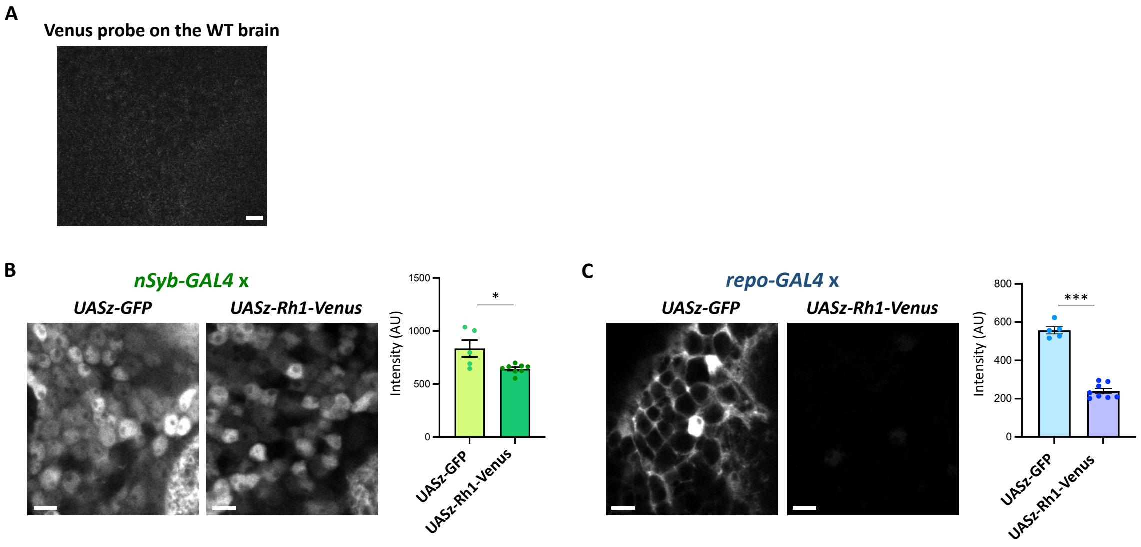


**Figure 5 – figure supplement 1. The *Rh1* reporter expression in neurons or in glia.**

(A) Negative control of the Venus smFISH. The Venus probes were hybridized onto the wild type brain and scanned with the identical setting to Figure 5E. (B-C) Expression of the *Rh1* (*UASz-Rh1-Venus*) or the control (*UASz-GFP*) reporter, driven by *nSyb-GAL4* (B) or by *repo-GAL4* (C). Sliced confocal images of the cortical regions adjacent to the antennal lobe are shown. Scale bars: 5 µm. The average fluorescent intensity is plotted. $N$ = 5 (*nSyb* > *UASz-GFP*), 8 (*nSyb* > *UASz-Rh1-Venus*), 5 (*repo* > *UASz-GFP*), 8 (*repo* > *UASz-Rh1-Venus*). *: $P < 0.05$, ***: $P < 0.001$, Mann-Whitney test of ranks.

Li, H., Janssens, J., De Waegeneer, M., Kolluru, S.S., Davie, K., Gardeux, V., Saelens, W., David, F.P.A., Brbić, M., Spanier, K., Leskovec, J., McLaughlin, C.N., Xie, Q., Jones, R.C., Brueckner, K., Shim, J., Tattikota, S.G., Schnorrer, F., Rust, K., Nystul, T.G., Carvalho-Santos, Z., Ribeiro, C., Pal, S., Mahadevaraju, S., Przytycka, T.M., Allen, A.M., Goodwin, S.F., Berry, C.W., Fuller, M.T., White-Cooper, H., Matunis, E.L., DiNardo, S., Galenza, A., O'Brien, L.E., Dow, J.A.T., FCA Consortium§, Jasper, H., Oliver, B., Perrimon, N., Deplancke, B., Quake, S.R., Luo, L., Aerts, S., Agarwal, D., Ahmed-Braimah, Y., Arbeitman, M., Ariss, M.M., Augsburger, J., Ayush, K., Baker, C.C., Banisch, T., Birker, K., Bodmer, R., Bolival, B., Brantley, S.E., Brill, J.A., Brown, N.C., Buehner, N.A., Cai, X.T., Cardoso-Figueiredo, R., Casares, F., Chang, A., Clandinin, T.R., Crasta, S., Desplan, C., Detweiler, A.M., Dhakan, D.B., Donà, E., Engert, S., Floc'hlay, S., George, N., González-Segarra, A.J., Groves, A.K., Gumbin, S., Guo, Y., Harris, D.E., Heifetz, Y., Holtz, S.L., Horns, F., Hudry, B., Hung, R.-J., Jan, Y.N., Jaszczak, J.S., Jefferis, G.S.X.E., Karkanias, J., Karr, T.L., Katheder, N.S., Kezos, J., Kim, A.A., Kim, S.K., Kockel, L., Konstantinides, N., Kornberg, T.B., Krause, H.M., Labott, A.T., Laturney, M., Lehmann, R., Leinwand, S., Li, J., Li, J.S.S., Li, Kai, Li, Ke, Li, L., Li, T., Litovchenko, M., Liu, H.-H., Liu, Y., Lu, T.-C., Manning, J., Mase, A., Matera-Vatnick, M., Matias, N.R., McDonough-Goldstein, C.E., McGeever, A., McLachlan, A.D., Moreno-Roman, P., Neff, N., Neville, M., Ngo, S., Nielsen, T., O'Brien, C.E., Osumi-Sutherland, D., Özel, M.N., Papatheodorou, I., Petkovic, M., Pilgrim, C., Pisco, A.O., Reisenman, C., Sanders, E.N., dos Santos, G., Scott, K., Sherlekar, A., Shiu, P., Sims, D., Sit, R.V., Slaidina, M., Smith, H.E., Sterne, G., Su, Y.-H., Sutton, D., Tamayo, M., Tan, M., Tastekin, I., Treiber, C., Vacek, D., Vogler, G., Waddell, S., Wang, W., Wilson, R.I., Wolfner, M.F., Wong, Y.-C.E., Xie, A., Xu, J., Yamamoto, S., Yan, J., Yao, Z., Yoda, K., Zhu, R., Zinzen, R.P., 2022. Fly Cell Atlas: A single-nucleus transcriptomic atlas of the adult fruit fly. Science 375, eabk2432. https://doi.org/10.1126/science.abk2432
